# Dpp Scaling is Not the Key to Robust Wing Patterning in *Drosophila*

**DOI:** 10.1101/2025.04.22.649574

**Authors:** Bhagyashree A. Ghag, Alexander W. Shingleton

## Abstract

Individuals within a population can vary dramatically in body and trait size in response to environmental factors such as developmental nutrition, temperature, and oxygen level, a phenomenon called phenotypic plasticity. Despite this variation, morphological pattern across the body and within traits are largely maintained, an example of developmental robustness. Consequently, the same trait can display both plastic and robust phenotypes. A fundamental yet unanswered question in developmental biology is: how do organs achieve plasticity in size while maintaining the robustness of pattern? This question is often answered using the *French flag model* of cell fate specification^1–4^. The model argues that cells receive positional information from a gradient of signaling molecules (morphogens) that diffuse from a single location in undifferentiated tissue. Specifically, the model argues that if the gradient is *dynamically scale invariant*, such that the gradient adjusts proportionally to the size of the tissue as it grows, then pattern is maintained regardless of the tissue size. However, while dynamic scale invariance has been demonstrated in several systems^5,6^, and explicitly^2^ or implicitly^7,8^ used to explain the robustness of pattern when tissue size varies due to environmental factors, this hypothesis has not yet been tested. We used the developing *Drosophila* wing as a model to test this hypothesis, under environmental conditions that reduce (low developmental oxygen level) and increase (low developmental temperature) adult wing size. We found that scale invariance of morphogen gradients is not observed under non-standard environmental conditions and is therefore not an explanation for robust patterning in the *Drosophila* wing with environmental variation in wing size.

## Results

### The Dpp signaling gradient is scale invariant under standard environmental conditions

Previous studies have suggested that the dynamic scale invariance of morphogen gradients, specifically the Decapentaplegic (Dpp)-signaling gradient, within the developing wing imaginal disc, accounts for the robustness of wing shape with wing size^2^. Dpp is secreted along the anterior-posterior axis of the wing imaginal disc to form a signaling gradient that defines the position of the longitudinal veins of the adult wing (**see Fig. S1A, A’, A”**). Earlier studies have demonstrated that the Dpp-signaling gradient is dynamically scale invariant, such that the decay length of the gradient (λ) is proportional to the width of the disc as the disc grows^5,9^. Thus, the relative position of the longitudinal veins, specified by the Dpp-signaling gradient, will be maintained regardless of disc and, hence, wing size (**see Fig. S1C, C’**).

To confirm the dynamic scale invariance of Dpp-signaling, we measured the relationship between the decay length of the Dpp signaling gradient, visualized by staining against pMAD, and disc size in discs ranging from 5 – 35×10^4^ μm^2^, collected from third instar larvae of different ages reared under standard environmental conditions (25°C, 21 kPa O_2_). After dissecting, staining, and imaging the discs, we fit an exponential function to the relationship between signal intensity and distance along the Dpp signaling gradient, normalizing intensity to lie between 0 and 1 (maximum intensity) and standardizing distance by multiplying length by the scaling factor *α*^−1/2^, where *α* is disc area (**Fig. 1A, B, C, D**). We then extracted the standardized decay length 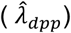 from the fitted exponential function (**Fig. 1D**). Consistent with previous studies, we found no significant relation between 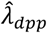 and disc size in larvae reared under standard laboratory conditions (LM, *R*^*2*^ = 0.001, *P* = 0.71), indicating that the Dpp-signaling gradient is scale invariant (**Fig. 2A, A’**). We also repeated our analysis using the methods of Wartlick *et al*. and again found that the Dpp-signaling gradient is scale invariant under standard laboratory conditions.

**Figure 1.**
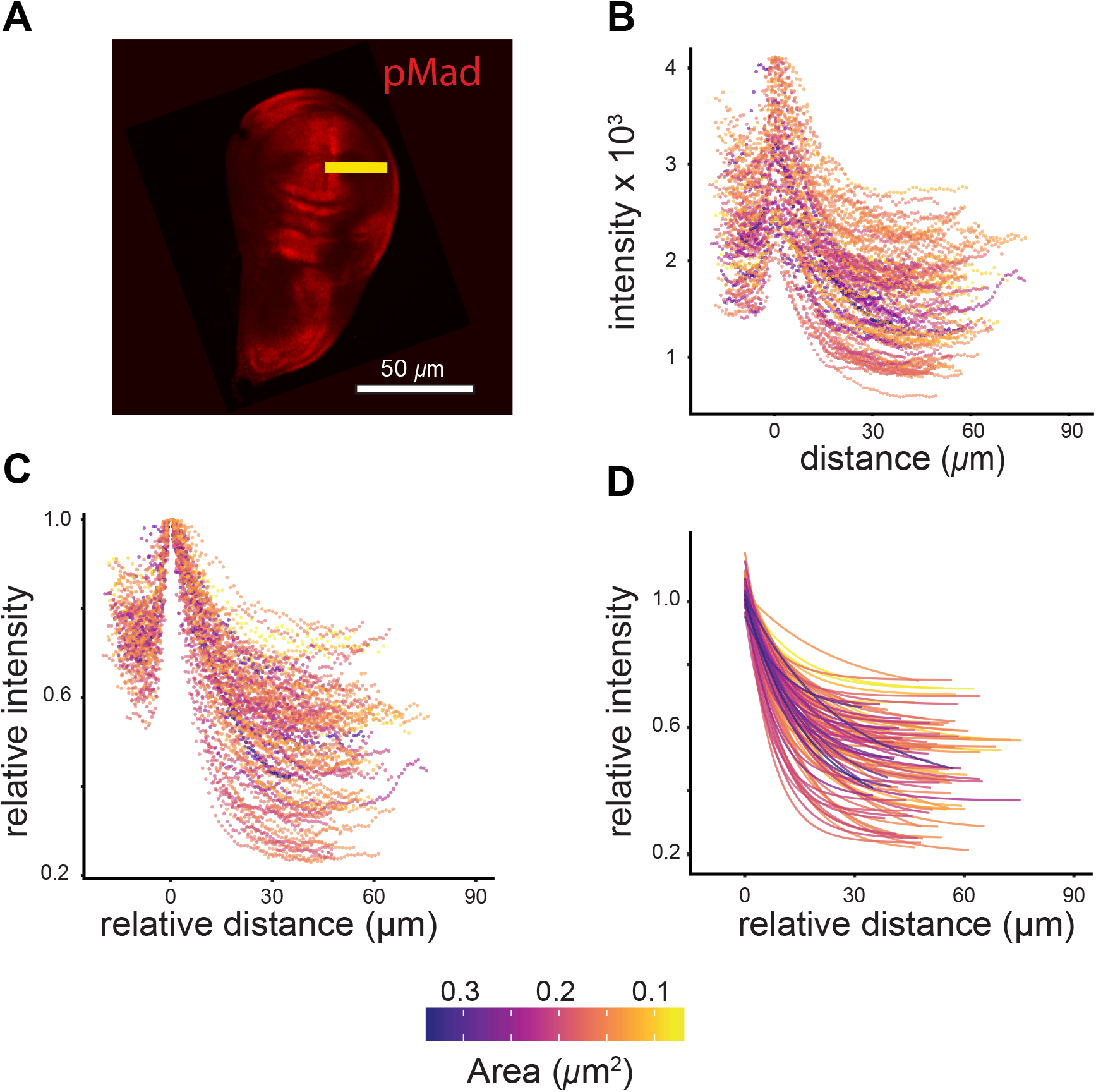
Analysing Dpp-signaling gradient. (**A**) A wing imaginal disc stained for Dpp-signaling gradient (pMad). Yellow bar is where intensity is read. (**B**) Raw intensity (*i*) against distance of pMad gradients in discs of different sizes. Gradients are centered at peak intensity. (**C**) Relative intensity of pMad gradients (normalized to *i*_*max*_ = 1) against relative distance (standardized by multiplying distance by the scaling factor *α*^−1/2^, where *α* is disc area) in discs of different sizes. (**D**) Exponential curves fit to intensity in (C) using 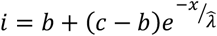, where *x* is distance, *b* is the lower asymptote, *c* is *i* when *x* = 0 and 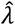 is standardized decay length of the signaling gradient (the position at which *i* has decreased to 1/*e* of *i*_*max*_ *=* 0.368).

**Figure 2.**
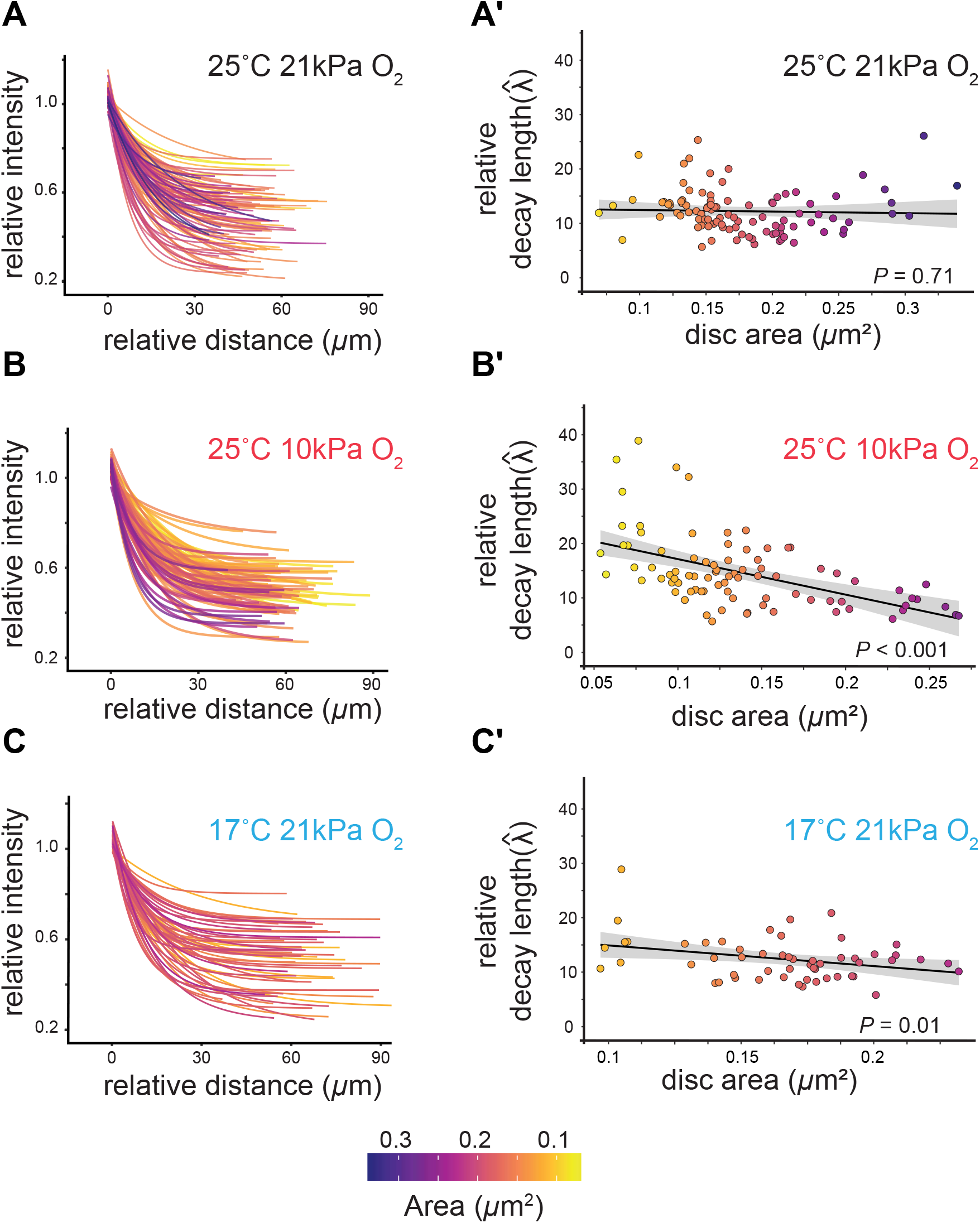
The Dpp signaling gradient is scale invarient under standard laboratory conditions while it underscales under non-standard laboratory conditions. (**A**) Exponential curve fitted to pMad gradients in discs of different sizes reared at standard conditions (25°C, 21 kPa O_2_) (same as Fig. 1D). (**A’**) the gradient is dynamically scale invariant under standard laboratory conditions, the gradient grows in proportion to the disc size (linear model: = *α*; standard conditions: *R*^*2*^ = 0.001, *P =* 0.71). (**B**) Exponential curve fitted to pMad gradients in discs of different sizes reared at low oxygen (25°C, 10 kPa O_2_). (**B’**) The gradient underscales with disc size, becoming progressively narrower as disc size increases (linear model: 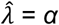; low oxygen: *R*^*2*^ = 0.275, *P* < 0.0001). (**C**) Exponential curve fitted to pMad gradients in discs of different sizes reared at low temperature (17°C, 21 kPa O_2_). (**C ‘**) The gradient underscales with disc size, becoming progressively narrower as disc size increases (linear model: = *α*; low temperature: *R*^*2*^ = 0.117, *P =* 0.0098)

### The Dpp signaling gradient is not scale invariant at low temperature and low oxygen levels

The dynamic scale invariance of the Dpp signaling gradient is an elegant explanation of the robustness of wing pattern with variation in wing size. However, it is only valid if the Dpp signaling gradient is dynamically scale invariant across the environmental conditions that generate variation in wing size. To test this, we measured the relationship between the decay length of the Dpp signaling gradient and disc size in third-instar larvae reared under conditions that reduce and increase wing size, compared to standard conditions. Specifically, we reared larvae at low oxygen (10 kPa O_2_, 25°C), which reduces wing size by 28 % (95% CI: 26-31%) compared to standard conditions (21 kPa O_2_, 25°C). We also reared larvae at low temperature (21 kPa O_2_, 17°C), which increases wing size by 29 % (95% CI: 27-32%) compared to standard conditions (**Fig. 3A**). Surprisingly, we found that at low temperature and low oxygen levels 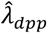 was not dynamically scale invariant (LM, *R*^*2*^ = 0.117, *P* = 0.0098 and LM, *R*^*2*^ = 0.275, *P <* 0.0001 respectively) (**Fig. 2B, B’, C, C’**). Specifically, in both conditions, the gradient underscales so that 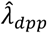 decreases, and the gradient becomes disproportionally narrower as the discs grow. Consequently, in discs from white-prepupae, just before metamorphosis, the Dpp signaling gradient is significantly narrower at low temperature and low oxygen compared to standard conditions (**see Fig. S2**). Thus, the dynamic scale invariance of the Dpp signaling gradient is not a general phenomenon but appears to be limited to standard environmental conditions.

**Figure 3.**
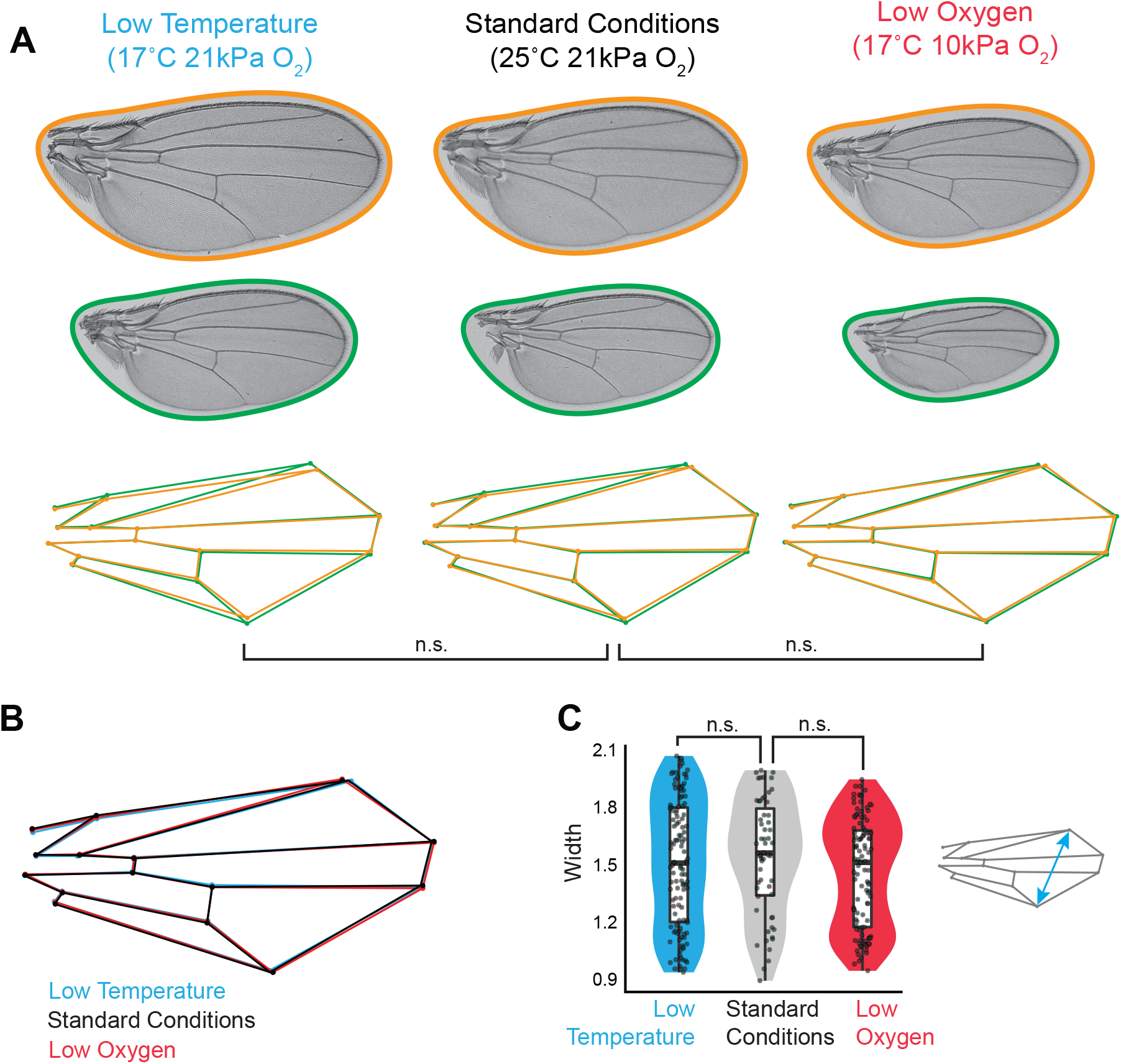
Wing pattern remains unchanged despite variations in Dpp-signaling gradient scaling across environments. (A) The largest and smallest wings from flies reared under each condition. Wireframes show allometric shape variation in each environmental condition, controlling for sex, between the largest (orange) and smallest (green) wings in each environmental condition. Neither temperature nor oxygen level interacts with the effect of centroid size on shape (Procrustes ANOVA, *F*_*temperature*size (1,173)*_ =1.273, *F*_*oxygen level*size (1,167)*_ = 0.546, *P* > 0.5 for both). (B) The effect of rearing environment on wing shape independent of size (non-allometric shape change). Wireframe shows mean wing shape in each environmental condition, controlling for centroid size and sex. (C) While the Dpp-signaling gradient underscales at low temperatures and oxygen levels, this is not reflected in a change in the width of the wing (cyan arrow on wireframe), controlling for centroid size and sex (Linear Model, *F*_*environment (2, 285)*_ *=* 0.9074 *P >* 0.4).

### Underscaling of the Dpp signaling gradient does not correlate with changes in wing pattern

If the scale invariance of the Dpp signaling gradient ensures wing-pattern robustness with wing-size variation under standard conditions, its absence under non-standard conditions should result in a corresponding lack of robustness as wing size varies. To test this, we used a morphometric analysis to assay the relationship between wing pattern and size within and between environmental conditions. Under standard environmental conditions (21 kPa O_2_, 25°C), there is a slight but significant change in wing shape with an increase in wing size, a phenomenon called allometric shape change (**Fig. 3A**). Nevertheless, size accounted for only 20% of the variation in shape among adult wings for flies reared under standard conditions, consistent with the general robustness of wing pattern with wing size.

Because the Dpp signaling gradient is not scale invariant at low temperature and low oxygen levels, we might expect there to be more substantial change in shape with size. This was not the case. The allometric component of wing shape variation – that is the change in wing shape that accompanies change in wing size – accounted for only 15% of total wing shape variation at low temperature levels, and only 6% at low oxygen levels (**Fig. 3A**). Further, the direction of allometric shape change – that is the way shape changes with size – was not different between standard and low temperature conditions (Procrustes ANOVA, *F*_*size*temp(1,167)*_ = 0.99, *P* = 0.863), nor between standard and low oxygen conditions (Procrustes ANOVA, *F*_*size*temp(1,180)*_ = 1.27, *P* = 0.230) (**Fig. 3A, B**). Thus, wing shape appears to be uniformly robust to variation in wing size, regardless of environmental condition.

Dpp defines the position of the longitudinal veins along the anterior-posterior axis of the wing^10,11^. Because the Dpp signaling gradient underscales at low oxygen and temperature and is disproportionally narrower at the end of the larval development (**Fig. 2A’, B’, C’**) we might expect the resulting adult wings to be disproportionally narrower compared to standard conditions. This was not the case. The width of the wing (the distance between landmarks 12 and 15, **Fig. S3**) was the same in flies reared under standard, low temperature and low oxygen conditions, when controlling for overall wing size (**Fig. 3C**). Indeed, the effect of low oxygen or low temperature on shape was very small. Comparing the wings from flies reared in standard versus low temperature conditions, temperature only accounted for 5% of the variation in shape, when controlling for the effects of sex and size. Similarly, oxygen level only accounted for 3% of the variation in shape when comparing wings from flies reared in standard versus low oxygen conditions, when controlling for the effect of sex and size. Collectively, therefore, while experimental changes in rearing environment generated 78% of the variation in size among wings, it accounted for only 8% of the variation in wing shape.

## Discussion

Dynamic scale invariance of morphogen gradients has been proposed as a key mechanism to generate robust patterning in organisms when body and organ size vary^2,7,8^. Indeed, it appears to be a central tenet in our understanding of morphogen action, supported by numerous studies demonstrating the scale invariance of several morphogen gradients. For example, the BMP activity gradient is scale-invariant along the dorsoventral axis in growing *Xenopus* embryos^12^, while β-catenin expression, part of the WNT-signaling pathway shows scale invariance in *Xenopus* embryos^13^. In *Drosophila*, Wartlick *et al*. demonstrated that the Dpp/BMP gradient in the wing imaginal disc is dynamically scale invariant along the anterior-posterior axis of the growing wing imaginal disc in *Drosophila* larvae^5^, and subsequent work demonstrated that this dynamic scale invariance is retained by downstream targets of Dpp-signaling^14^.

Critically, however, all the above-mentioned studies were conducted under standard laboratory conditions chosen to optimize organismal growth and reduce body and organ size variation. Such conditions are far removed from the environmental variation experienced by growing animals in nature, which generates considerable variation in size within a genotype. Conceptually, there are four possible models of the relationship between the decay length of the morphogen gradient 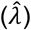 and disc size (*a*) when environments vary. The first (Model 1) is that the relationship is dynamically scale-invariant and the same regardless of environmental conditions (**Fig. 4A**). Under this model, we would expect final pattern to be maintained within and between environments. The second (Model 2) is that the relationship is scale-invariant but different between environmental conditions (**Fig. 4B**). Under this model, we would expect final pattern to be maintained within environments but be different between environments. The third (Model 3) is that the relationship is not scale invariant, but the decay length of the morphogen gradients converge at the final disc size, referred to as static scale invariance (**Fig. 4C**). Under this model, we would expect the final pattern to be maintained between environments but be different within environments. The fourth model (Model 4) is that there is neither dynamic nor static scale invariance within or between all environments (**Fig. 4D**). Under this model we would expect the final pattern to not be robust but vary with size both within and between environments.

**Figure 4.**
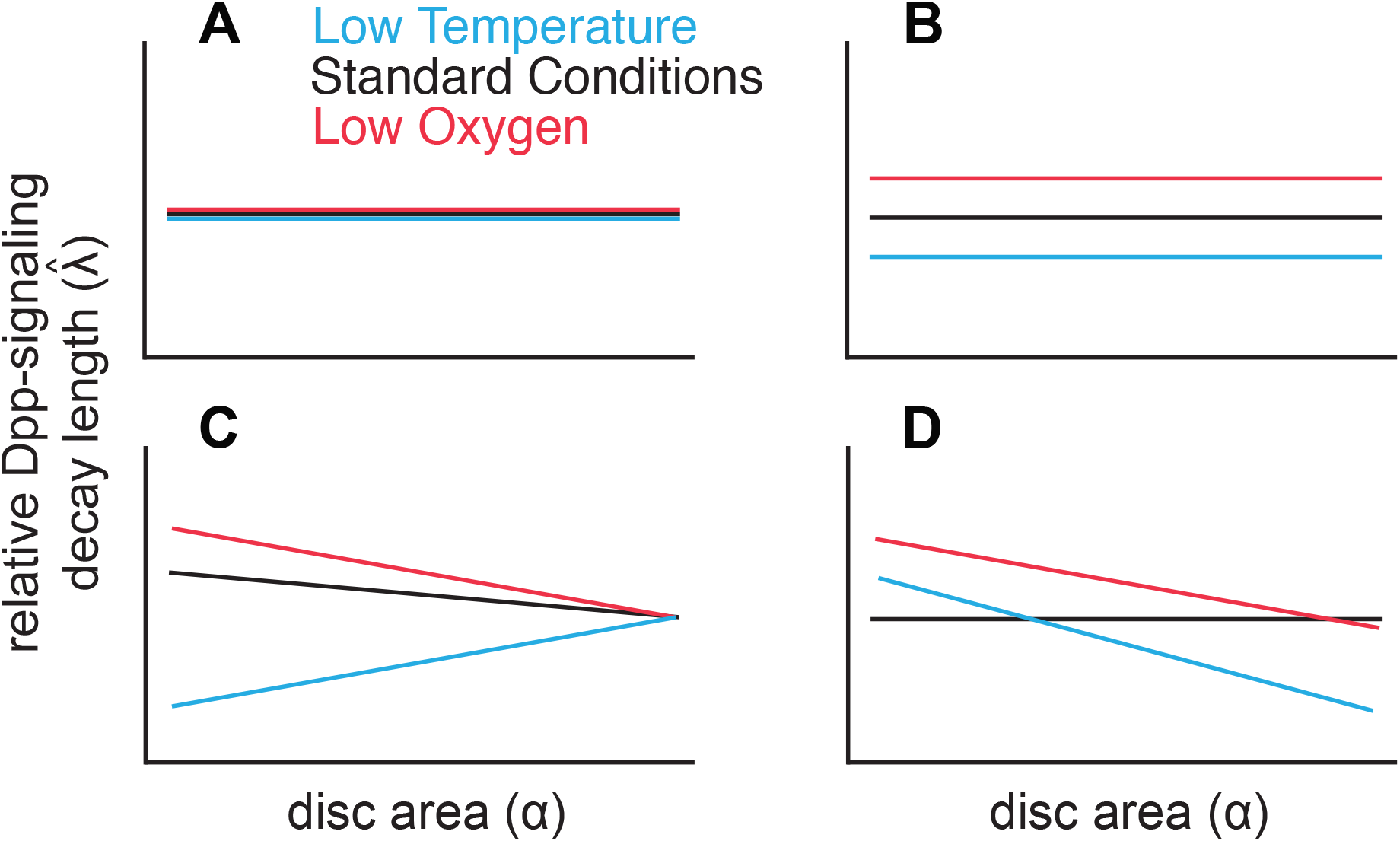
Hypothetical relationships between standardized decay length, 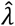, of the Dpp-signaling gradient and disc size, α, under different environmental conditions. (A) Dynamic scale invariance across environments: 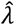 is independent of *α* and identical across all environmental conditions; (B) Environment-dependent dynamic scale invariance: 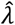 is independent of *α*, but varies between environments; (C) Static scale invariance: 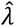 varies with *α* in all or some of the environments, but converges on the same value at the end of ontogeny.(D) No scale invariance: 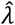 varies with *α* in all or some of the environments and does not converge on the same value at the end of ontogeny.

Data from our analysis of the Dpp signaling gradient align with Model 4. Based on this model, if scale invariance of the Dpp signaling gradient were responsible for maintaining robustness of wing shape, we would anticipate corresponding changes in wing shape with variations in size, both within and between environmental conditions. However, this is not the case. Our data indicate that while wing shape does change marginally with wing size (**Fig. 3A**), consistent with earlier papers^15-17^, this change in shape is the same across environmental conditions, so it cannot be explained by the dynamic scale invariance of the Dpp signaling gradient. Further, our data indicate that the Dpp signaling gradient is not statically scale invariant (Model 3) (**see Fig. S2**). While the Dpp signaling gradient is disproportionally narrow at low oxygen and low temperature conditions by the end of larval development, this is not reflected by a narrowing of the adult wings (**Fig. 3C**). In general, therefore, there is a lack of correspondence between dynamic and static scaling of the Dpp-signaling gradient and changes in wing shape with wing size within and between environmental conditions.

The observed change in scaling of the Dpp-signaling gradient under non-standard environmental conditions is not due to the inadequacy of our experimental methods to detect dynamic scale invariance. First, the null hypothesis of our statistical analysis is dynamic scale invariance; that is, that there is no statistical relationship between between 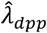 and *α*. Consequently, our observation of changes in Dpp scaling under non-standard rearing conditions cannot be due to insufficient statistical power to reject the null hypothesis. Indeed, a *post hoc* power analysis indicated that our study had 80% power to detect an effect of environment on the relationship between 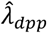 and *α* as small as *R*^*2*^ = 0.054 (the observed *R*^*2*^ was 0.22). Second, our methods were sufficient to detect dynamic scale invariance under standard laboratory conditions. Finally, we repeated our analysis using the methods of Wartlick *et al* and found the results unchanged. Thus, our findings are unlikely due to our experimental or analytical methods.

If scale invariance of the Dpp-signaling gradient does not account for the robustness of wing pattern with wing size in *Drosophila*, how then is this robustness conferred? It is possible that scale invariance is attained further down the Dpp-signaling cascade. Previous research established the scale invariance of brinker (Brk), daughters against dpp (dad), spalt (Sal), optomotor-blind (omb), pentagone (pent), and delta (Dl) ^14^. The products of these genes show distinct spatial patterns of expression, be they gradients (e.g., Brk, which forms a gradient the inverse of Dpp) or discrete expression patterns (e.g., Dl, which marks the positions of cells that become L1,3,4,5). These spatial patterns could be dynamically or statically scale invariant. Exploring this possibility represents the next phase of our research project, promising to unravel further intricacies of developmental patterning and shed light on the mechanisms underlying robustness in diverse environmental contexts.

In summary, our findings challenge the hypothesis that morphogen scale invariance is necessary for robust patterning. Despite the Dpp gradient failing to scale dynamically or statically under a range of environmental conditions, wing shape remains remarkably stable. This suggests that an alternative regulatory mechanism buffers developmental variation in the expression of patterning genes when wing size varies across environments. Future research should explore targets of Dpp signaling, such as Brk and pent, to determine where downstream of the Dpp-signaling gradient robustness is conferred. Our work highlights the need to reconsider the role of morphogen scaling in developmental robustness and suggests that compensatory mechanisms beyond gradient scaling maintain pattern fidelity.

## Supporting information

STAR Material and Methods

## Supplemental information

STAR Material and Methods, Supplemental Figures S1-S3

