## Supplementary material for "Dpp Scaling is Not the Key to Robust Wing Patterning in *Drosophila*": STAR Material and Methods

### STAR methods

### KEY RESOURCES TABLE

| REAGENT/RESOURCES | SOURCES | IDENTIFIER |
|---|---|---|
| Antibodies | | |
| Anti-Smad3 (phospho S423 + S425) antibody [EP823Y] (1:50) | Abcam | Cat:NC9026662 |
| Goat anti-Rabbit IgG (H+L) Secondary Antibody, Alexa Fluor™ Plus 594, Invitrogen™ (1:200) | Fisher Scientific | Cat: PIA32740 |
| Reagents | | |
| PBS | Fisher Scientific International Inc. | Cat: BP3994 |
| Triton X | RPI | Cat: 111036 |
| i-Block | ThermoFisher Scientific | Cat: T2015 |
| Normal Goat Serum (NGS) | Abcam | Cat: ab7481 |
| Vectashield | Vector laboratories | Cat: H-1200-10 |
| Paraformaldehyde (PFA) | Fisher Scientific Company | Cat: T-353 |
| Dimethyl Hydantoin Formaldehyde (DMHF) | Entomopraxis | Cat: A9001 |
| Software | | |
| R | R version 4.3.2 (2023-10-31 ucrt) | |
| JMP | JMP Pro 17.2.0 | |
| ImageJ | v1.54k | |
| MorphoJ | 1.08.02 | |
| Other | | |

| |
|---|
| Olympus Optical FV-500 laser scanning confocal microscope |
| Leica DM6000B epifluorescence microscope |

### EXPERIMENTAL MODEL AND SUBJECT DETAILS

All experiments used $w^{118}$ flies from the Vienna *Drosophila* RNAi Center (Stock # 60,000). Unless otherwise stated, flies were reared at low density (50-100 larvae per vial) on cornmeal-molasses medium[18] under standard environmental conditions (25°C, 21% $O_2$) and constant light.

### METHOD DETAILS

All data (images and measurements) and the scripts used to analyze them are provided on Dryad and as supplemental information (Script S1-2, Data S1-4). All model fitting was conducted in *R*.

*Dpp-Signaling Assay: Fly Rearing and Experimental Conditions*

Females were allowed to oviposit for 24 hours on standard cornmeal-molasses medium, yielding ~50–100 eggs per vial. Eggs/larvae were then maintained under standard (25°C, 21 kPa $O_2$), low temperature (17°C, 21 kPa $O_2$), or low oxygen (17°C, 10 kPa $O_2$) conditions. Once the oldest larva in each vial reached the white pre-pupae stage, all third instar larvae were dissected for wing disc immunostaining. This approach captured a range of developmental stages and corresponding wing disc sizes. Larvae were collected from 30 vials per condition.

*Dpp-Signaling Assay: Wing Disc Immunostaining*

Wing imaginal discs were dissected in ice-cold PBS, fixed in 4% PFA for 25 minutes, and permeabilized in PBT (0.3% Triton X in PBS) for 1 hour. They were then blocked in 2% normal goat serum in BBT (0.2% I-Block in PBT) for 1 hour at room temperature. Next, the samples were incubated with anti-pMad primary antibody (1:50 in BBT/NGS) for 2 hours with gentle rocking, followed by four 15-minute washes in PBT. After an additional 1-hour block in BBT/NGS, they were incubated overnight at 4°C with Alexa Fluor™ Plus 594-conjugated secondary antibody (1:200 in BBT/NGS), protected from light. The following day, the discs were washed four times for 15 minutes each in PBT, mounted in Vectashield, and stored at 4°C in the dark until confocal microscopy analysis.

*Dpp-Signaling Assay: Microscopy and Image Acquisition*

Immunostained wing imaginal discs were imaged on an Olympus FV-500 confocal microscope. Z-stacks were acquired at 1.5 μm intervals with a 10X objective. The number of z-stacks varied depending on the thickness of each disc, but typically, around 20-30 stacks were captured per disc. The upper and lower limits of the stacks were set to traverse the entire depth of the pouch. Identical microscope and camera settings were applied for all image capture, ensuring consistency across standard, low temperature (17°C), and low oxygen (10 kPa $O_2$) conditions. Maximum intensity projections were then generated in ImageJ for gradient analysis[5].

*Morphometric Analysis: Fly Rearing, Experimental Conditions, Microscopy and Image Acquisition*

Flies were reared as in the Dpp-signaling assay: After a 24-hour oviposition at 25°C and 21kPa $O_2$, eggs were transferred to one of three conditions (standard, low temperature, or low oxygen) and reared to the pupal stage. Pupae were individually collected in Eppendorf tubes under their respective conditions. Upon eclosion, adults were preserved in 70% ethanol, their right wings dissected, mounted in DMHF, and imaged using a Leica DM6000B20. Flies were collected from 20 vials per condition.

### QUANTIFICATION AND STATISTICAL ANALYSIS

*Dpp-Signaling Assay*

The Dpp-signaling gradient was assayed as the intensity profile of pMad staining in the posterior compartment of the wing imaginal disc, down a strip extending from peak signal at the anterior-posterior boundary to the posterior edge of the disc [9] (Fig. 1A). Intensity ($i$) was normalized by setting maximum intensity to 1 and distance was standardized relative to wing disc size by multiplying by $\alpha^{1/2}$, where α is disc area. We then used a non-linear model to fit the exponential curve $i = b + (c - b)e^{-z/\hat{\lambda}}$ , to the data, where $x$ is distance, $b$ is the lower asymptote, $c$ is $i$ when $x$ = 0 and $\hat{\lambda}$ is the standardized decay length of the gradient profile (the position at which $i$ has decreased to 1/$e$ of $i_{max}$ = 0.368). If the Dpp-signaling gradient were scale invariant, is constant regardless of disc size. To test this, we regressed $\hat{\lambda}$ against disc area α, using an ordinary least square regression model.

*Morphometric Analysis*

The coordinates of 15 landmarks across each wing were captured using the *Fly_Wing_15lmk* plugin (kindly provided by Ian Dworkin) in ImageJ[19]. We used MorphoJ to conduct a global Procrustes transformation on all wings and visually inspected the scatterplots of superimposed Procrustes coordinates for gross outliers. Procrustes-transformed coordinates and centroid size were then imported into *R* and analyzed using the *geomorph* package. To test the effect of size, sex and environment on wing shape we analysed the data using a Procrustes ANOVA by fitting the model $\boldsymbol{P}_{\mathrm{ijkl}} = C_i+S_j+E_k+C{\cdot}S_{ij}+C{\cdot}E_{ik}+S{\cdot}E_{jk}+C{\cdot}S{\cdot}E_{ijk}+\varepsilon_{ijkl}$, where $\boldsymbol{P}$ is the matrix of Procrustes-transformed landmark coordinates, $C$ is wing centroid size, $S$ is sex, $E$ is environmental condition and ε is error (subscripts are levels within each parameter). Non-significant factors and interactions were removed from the model and the data were re-analysed using the simpler model. Within each environmental condition we fit the model $\boldsymbol{P}_{\mathrm{ijk}} = C_i+S_j+\varepsilon_{ijk}$ and calculated the $R^2$ of effect of size on shape as the proportion of total shape variance explained by size. To test the effect of the environment on wing width, we fit the model $W_{ijk}=S_i+E_j+S{\cdot}E_{ij}+\varepsilon_{ijk}$, where $W$ is the Procrustes-transformed wing width (distance between landmarks 12 and 15, Fig. S3). As before, non-significant factors and interactions were removed from the model and the data were re-analysed using the simpler model.
