## Supplemental Figures for "Dpp Scaling is Not the Key to Robust Wing Patterning in *Drosophila*"

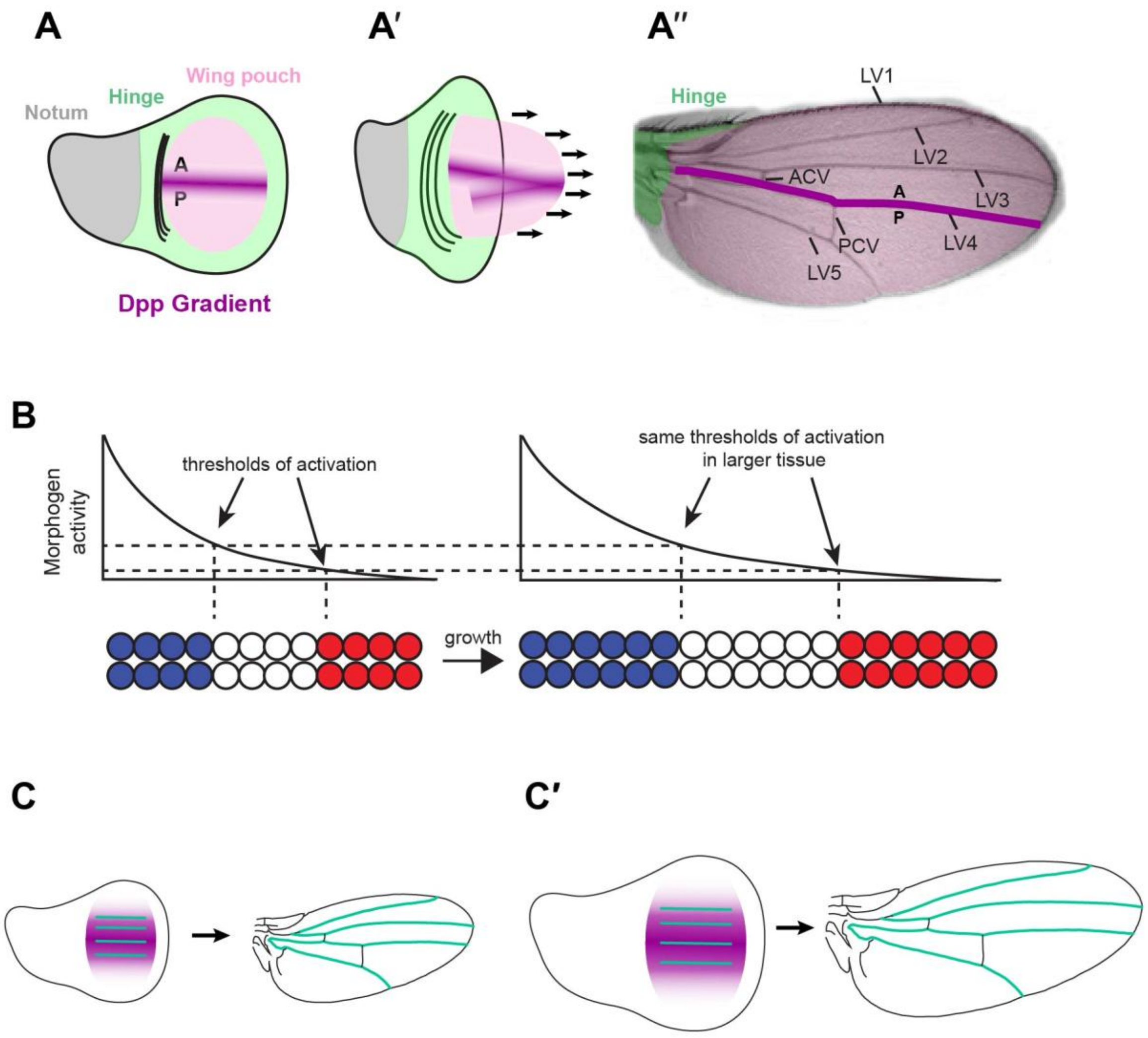


Figure S1: A) The wing imaginal disc is divided into, hinge, notum, wing pouch. The wing blade is divided into anterior-posterior (A-P) by the Dpp gradient. A') The wing disc everts (arrows) so that the Dpp gradient defines the position of the latitudinal veins (L1-5). A") Pattern of the adult wing, showing the A-P axis.B) The French flag model-The threshold responses to a signaling gradient provides positional information. If the gradient is scale invariant as tissues grow, the same pattern will be generated across tissues of different sizes. C) If the Dpp gradient is dynamically scale invariant then the same vein pattern will generated in small and (C′) large wings.

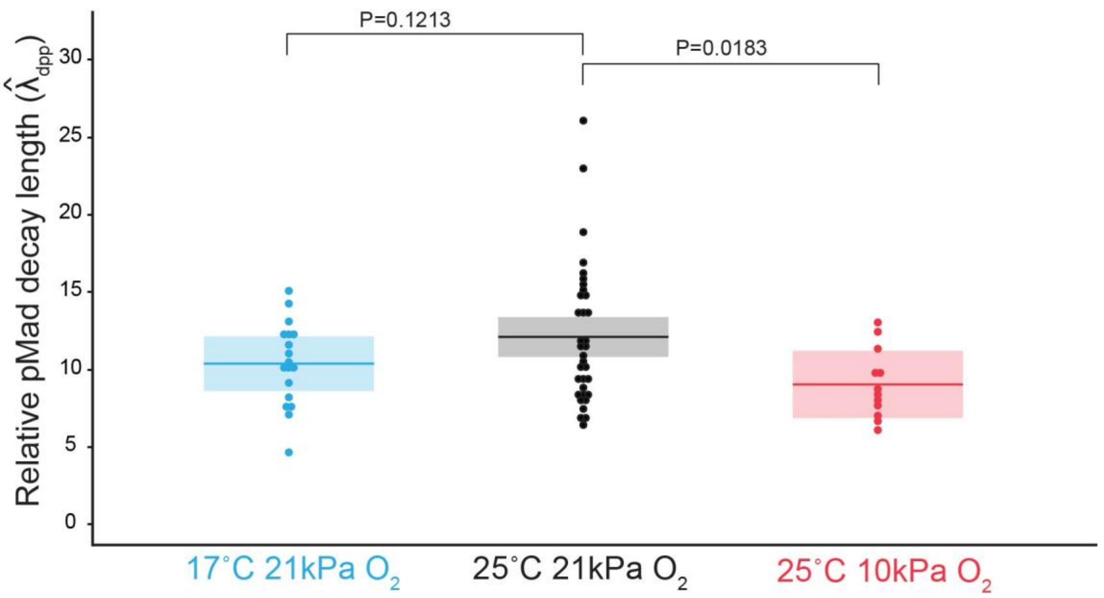


Figure S2: Dpp signaling gradient is not statistically scale invariant, related to Figure.2. Relative decay length ($\hat{\lambda}_{dpp}$ ) of Dpp signaling gradient at the end of the third larval instar in standard conditions, low temperature condtions, and low oxygen conditions. The boxes represent the mean ± 95% confidence interval. *P*-values are from unpaired two-tailed t-tests comparing each condition to the standard.

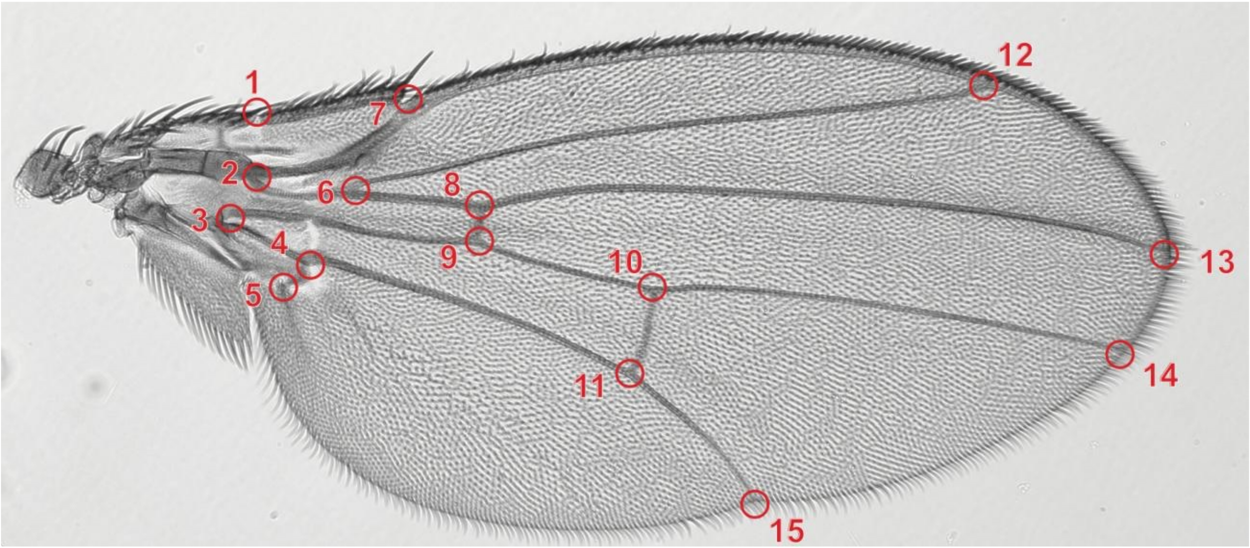


Figure S3: Location of the 15 landmarks used in the morphological analysis.
